# Social and individual learners use different pathways to success in an ant mini-society

**DOI:** 10.1101/2022.07.01.498419

**Authors:** Simone M Glaser, Christoph Grüter

**Affiliations:** Institute of Organismic and Molecular Evolutionary Biology, Johannes-Gutenberg University of Mainz, Germany; School of Biological Sciences, University of Bristol, 24 Tyndall Avenue, BS8 1TQ Bristol, UK

**Keywords:** recruitment, social learning, tandem running, *Temnothorax nylanderi*

## Abstract

Animals can acquire information through individual learning or by copying others. Simulations suggest that social learning is expected to lead to better rewards, but experimental studies confirming this remain scarce. We tested how a well-known form of social learning in ants, tandem running, affects individual foraging success of *Temnothroax nylanderi* foragers in controlled laboratory experiments. We manipulated the number and the variability of food sources and assessed the foraging choices of ants searching individually (*i*.*e*. scouts) or using social learning (*i*.*e*. recruits). We found that social learners indeed discovered better food sources than individual learners, but only in rich environments. However, social learners collected less food (∼60% fewer foraging trips) than scouts during our trials. Interestingly, individual learners improved their success over time by switching food sources more frequently than social learners. These experimental findings highlight that the relative value of social and individual learning in an ant society depend on the foraging environment and show different temporal dynamics. The ability of individual learners to exploit profitable food sources through a strategy of food source switching, while avoiding the opportunity costs of social learning can help explain why many social insects, especially those living in small colonies, do not use communication in foraging.

## Introduction

Animals need to acquire and update information about their environment to make adaptive decisions, for example, when searching for food sources, nesting sites and mating opportunities or to avoid predators (Boyd and Richerson 1995; Brown and Laland 2003; Danchin et al. 2004; Dall et al. 2005; Horner et al. 2010; Slagsvold and Wiebe 2011; Dawson and Chittka 2014). Individuals can collect new information via individual (trial-and-error) learning or they can use social learning, *i*.*e*. learning by observing or interacting with other individuals or their products (Laland 2004; Galef and Laland 2005; Rendell et al. 2010; Shettleworth 2010; Heyes 2012). If an individual already has acquired information, it can rely on its memory and use this “private information” (Kendal et al. 2005; Grüter and Leadbeater 2014).

Social information is often considered to be a low-cost option, whereas individual learning might be more costly to collect but is also more accurate (Laland 2004; Rieucau and Giraldeau 2011). Private information, on the other hand, is immediately available, but is prone to being outdated (e.g. Kendal et al. 2005; Rendell et al. 2010; Grüter and Leadbeater 2014; Smolla et al. 2016). However, still little is known about the relative payoffs of the different strategies, and how these depend on the ecology of a species. Rendell et al. (2010) used a simulated computer tournament to test the success of different information-use strategies and found that social learning tended to be more successful than individual learning because “demonstrators” use and, therefore, display the most productive behaviour in their repertoire, thereby “filtering” information for observers. However, experimental evidence showing that social learning leads to better rewards than individual learning remains scarce (Rendell et al. 2010; but see e.g. Thornton and McAuliffe 2006).

Eusocial insects are well-known for relying on social information during foraging or colony migration (von Frisch 1967; Leadbeater & Chittka 2007; Grüter & Leadbeater 2014). Honeybees (*Apis* spp.), for example, use the waggle dance communication (von Frisch 1967) and many ants, termites and stingless bees rely on trail pheromones (Hölldobler and Wilson 2009; Hrncir and Maia-Silva 2013; Czaczkes et al. 2015; Grüter 2020). Empirical studies confirm that ant or honeybee foragers often only share information with nestmates when the food source is of high quality (von Frisch 1967; reviewed in Grüter and Czaczkes 2019), which leads to information filtering as described in Rendell et al. (2010). As predicted, there is evidence that honeybees following waggle dances find better food sources than scouts using an individual learning strategy (Seeley 1983; Seeley and Visscher 1988), but dance recruits also required more time to locate a food source, highlighting that social learning has considerable time and opportunity costs (Seeley 1983; Seeley and Visscher 1988; Dechaume-Moncharmont et al. 2005; Franks and Richardson 2006; Grüter et al. 2010; I’Anson Price et al. 2019).

In a cooperative society such as an insect colony, an information-use strategy that increases rewards for the individual might not necessarily be beneficial for colony success (Grüter and Leadbeater 2014). For example, colonies that rely strongly on social learning could be worse off if social learning has considerable time costs (Dechaume-Moncharmont et al. 2005; I’Anson Price et al. 2019). Theoretical models of honeybee foraging suggest that both the benefits and the costs of social information depend on food source distribution (Dornhaus et al. 2006; Beekman and Lew 2007; Schürch and Grüter 2014; I’Anson Price et al. 2019; Goy et al. 2021). When food sources are easy to find, individual learning is often more efficient. This could explain why many social insect species do not use communication about food source locations. Bumblebees, many stingless bees and many ants with small colony sizes, for example, forage largely solitarily (foragers may use social cues at food sources to make foraging decisions) (Beckers et al. 1989; Leadbeater and Chittka 2005; Worden and Papaj 2005; Dunlap et al. 2016; Grüter 2020). Thus, learning socially about where to find food might often not improve colony foraging success.

One reason for our limited understanding of the adaptive value of social learning in social insects is the difficulty to distinguish between social and private information use (Grüter and Farina 2009; Czaczkes et al. 2016). For example, bees might follow a waggle dance, but subsequently use private information to fly to a different food source after leaving the nest (Grüter et al. 2008, 2013; Menzel et al. 2011) and ants walking on a trail marked by pheromone might rely on memory rather than chemicals or a combination of both when making decisions (Grüter et al. 2011; Czaczkes et al. 2011, 2015).

Tandem running is an ideal behaviour to study the outcome of social learning because social learners always pair up with another ant (Wilson 1959; Möglich et al. 1974; Franks and Richardson 2006; Franklin 2014). This form of communication is used in many ant species with small colony sizes and involves an experienced leader (tutor) and a naïve follower (henceforth also called recruit or social learner) (Möglich et al. 1974; Franks and Richardson 2006; Richardson et al. 2007; Franklin 2014; Glaser and Grüter 2018; Grüter et al. 2018; Silva et al. 2021). A tandem run starts when a successful forager returns to their nest and attracts a potential follower, often by producing an attractive pheromone (“tandem calling”) (Möglich et al. 1974). After learning the route to the food source while being guided by a leader (Sasaki et al. 2020), the follower can herself become a tandem leader and guide other ants to the food source.

We studied the common European ant *Temnothorax nylanderi* as a model system and studied individual foraging success in controlled environments that differed in the number and variability of available food sources. We tracked individually marked social and individual learners over repeated foraging trips, analysed their foraging decisions and recorded the outcome of each trip. Our first (*i*) aim was to quantify tandem running in different environments. Secondly (*ii*), we tested the prediction that social learners (recruits) find food sources of higher quality than individual learners (scouts). We also tested (*iii*) if individuals can improve rewards over successive trips by abandoning food sources that are suboptimal. Previous research suggests that social learners experience greater time costs as they wait inside the nest for information (Seeley 1983; Seeley and Visscher 1988; Dechaume-Moncharmont et al. 2005; Goy et al. 2021), thus we (*iv*) predicted that social learners perform fewer foraging trips than individual learners during our trials.

## Material & Methods

### Study site and study species

Thirty colonies of *Temnothorax nylanderi* were collected in the Lenneberger forest near Mainz, Germany. Colonies were kept in climate chambers at 25°C with a 12:12 h light/dark cycle. The colonies lived in nests that consisted of two microscope slides (50 × 10 × 3 mm) and between the two slides was a plexiglass slide with an oval cavity and an opening as a nest entrance. The nest was placed in a larger three-chamber-box (100 × 100 × 30 mm) and to prevent the ants from escaping, the walls were coated with paraffin oil. Colonies were provided with an *ad libitum* water source and fed twice a week with honey and cricket. All colonies had a reproductive queen, brood and a mean colony size (adult workers) of 77.0 ± 34.8 (± StDev).

### Experimental set-up and procedure

All experiments were conducted in the same climate chamber. Colonies were starved for 10 days before each trial to guarantee foraging motivation. After seven days of starvation the nests were placed in a foraging arena (78 × 56 × 18 cm), which allowed ants to get used to the test environment. The floor of the arena was covered with white paper to improve contrast for filming. The paper was changed after each trial with a colony to remove potential chemical traces (but note that *Temnothorax* foragers are not known to follow pheromone trails deposited on the ground to food sources; Möglich et al. 1974; Franklin 2014). To prevent ants from escaping, the walls of the arenas were covered with Fluon.

Fifteen colonies were tested in four conditions each: with two and ten food sources of variable (0.1M and 1.0M sucrose solution) or constant (1.0M) quality. The nest was always positioned in the middle of the arena. The testing order was randomized for each colony. On an experiment day, either two or ten droplets of sucrose solution (ca. 50 μl) were provided, which was enough to last for the duration of a trial (the crop load of an individual ant is ∼0.15 μl, unpublished measurements) (Fig. S1). Ants of the species *T. nylanderi* typically forage less than 50 cm from the nest (Heinze et al. 1996). Therefore, sugar droplets were positioned 20 cm from the nest entrance with equal distances between food sources. In the case of ten variable food sources, they were positioned in alternating order around the nest, *i*.*e*. a 1M food source had a 0.1M food source on either side and *vice versa* (Fig. S1). After the first forager discovered a food source, video recording (SONY HDR-CX200) started for 90 min. When scouts (*i*.*e*. individual learners = ants that left the nest alone to locate their first food source) or tandem followers (social learners, recruits) reached the food source, they were individually marked with a colour dot on their abdomen (POSCA, Mitsubishi Pencil Co., UK). This allowed us to follow the foraging success of individually marked foragers over several foraging trips. If possible, four scouts and four followers were marked per trial. Thus, ants were classified as either individual or social learners based on how they discovered their first food source. After every trial, colonies were fed for four days before being starved again to be tested in a different setup. Thus, the trial period was 14 days and included 10 days without any food sources in their environment.

In additional experiments, we explored time costs associated with using social or individual learning by measuring if followers require more or less time than scouts to discover their first food source after leaving the nest. Fifteen different colonies were provided with either two or ten sugar droplets of good quality (1.0 M) sucrose solution. The video recording started after a scout left the nest. We filmed until colonies performed two tandem runs to compare the nest-to-food time of scouts and tandem followers.

### Data collection

Video recordings were analysed with the VLC media player V5. We recorded which of the following foraging strategies our marked ants used to locate food sources: scouting (individual learning), returning to food alone (private information user), leading a tandem run (private information user) or following a tandem run (social learning) to food. We also noted if our focal ants switched food sources and whether they visited a good or a bad food source. We calculated the success rate of tandem runs and the probability to perform a tandem run for each visit. A tandem run was considered successful (*1*) if the pair reached the food source together or (*2*) if a follower was guided to within 1 cm from the food source and discovered the food source afterwards (similar to Glaser and Grüter 2018). In the second experiment, we recorded the time that scouts and tandem followers needed from the nest to a food source.

### Statistical analysis

All statistical tests were done in R 3.4.2 (Team 2017). For normally-distributed response variables we used linear mixed-effect models (LME). For response variables with a binomial or Poisson distribution, we used generalized linear mixed-effect models (GLMM) (Zuur et al. 2009). We tested if models with a Poisson distribution were overdispersed (package blmeco; Barry et al. 2003). If necessary, variables were transformed (square root or boxcox transformation) to achieve normality and then analysed with an LME. Colony ID and ant ID were used as random effects to control for the non-independence of data points from the same colony and the same ant (Zuur et al. 2009).

Different questions were addressed by inclusion of different fixed effects: *foraging setup* (two-variable, two-constant, ten-variable and ten-constant food sources), *strategy* (scout, leader, follower and private information user), *food quality* (good or bad, used when analysing food source switching) and *experience*. The effect of *foraging experience* was analysed as the number of food source visits during a trial. Most of our models (*i*.*e*. predictions) had just one predictor, but we had a few models with two predictors. In these models, we tested the interaction between the two variables with Likelihood Ratio tests (LRT). For instance, previous research has shown that foraging experience affects tandem probability (Glaser and Grüter 2018), but it is not yet known if this effect depends on food source quality. Therefore, the model contained *foraging setup* as a second predictor to test for a possible interaction between *experience* and *foraging setup* on the probability to perform tandem runs (Zuur et al. 2009). An interaction was removed from the model if it was not significant (Zuur et al. 2009).

As response variables, we used tandem success rate (successful vs. unsuccessful, binomial GLMM), the probability of tandem runs (yes vs. no, binomial GLMM), the probability of food switches (switched vs. stayed, binomial GLMM), quality of food source discovered (high vs. low, binomial GLMM), the total number of foraging trips (Poisson GLMM) and the time that scouts and tandem followers needed to reach a food source (Gaussian LME). We used a boxcox transformation to achieve normality to test whether the duration from nest to food source depended on the setup (lambda = -0,4).

## Results

### (i) Tandem success rate and probability to perform tandem runs

The success rate of tandem runs (total N = 747) was higher in environments with ten food sources (Fig. 1) (GLMM: variable: two vs. ten: z = 4.341, p < 0.001; constant: two vs. ten: z = 2.435, p < 0.001). The success rate did not depend on whether food sources had the same or different quality (two: 55.5% of 290 vs. ten: 75.5% of 457; GLMM: two: variable vs. constant: z = 1.535, p = 0.125; ten: variable vs. constant: z = 0.211, p = 0.832), suggesting that tandem runs to bad food sources were not less successful. Unexpectedly, the probability to perform a tandem run was significantly lower when there were more food sources (Fig. 2) (two: 53.4% vs. ten: 31.7%; GLMM: two: variable vs. constant: z = 0.638, p = 0.523; ten: variable vs. constant: z = 0.252, p = 0.801; variable: two vs. ten: z = -3.377, p < 0.001; constant: two vs. ten: z = 3.725, p < 0.001). The probability that ants performed a tandem run increased with their experience (measured as the number of previous visits) (GLMM: two vs. ten: z = 0.369, p = 0.712; visit: z = 3.188, p = 0.001; interaction: z = 0.257, p < 0.001). The significant interaction between food source number and visit number indicates that the probability of tandem runs increased more with experience when there were two food sources (based on visual inspection after plotting the results). Furthermore, the probability to start a tandem run was significantly higher after visiting a good food source (Fig. 3A) (GLMM: two: z = 3.669, p < 0.001; ten: z = 4.989, p < 0.001).

**Fig. 1.**
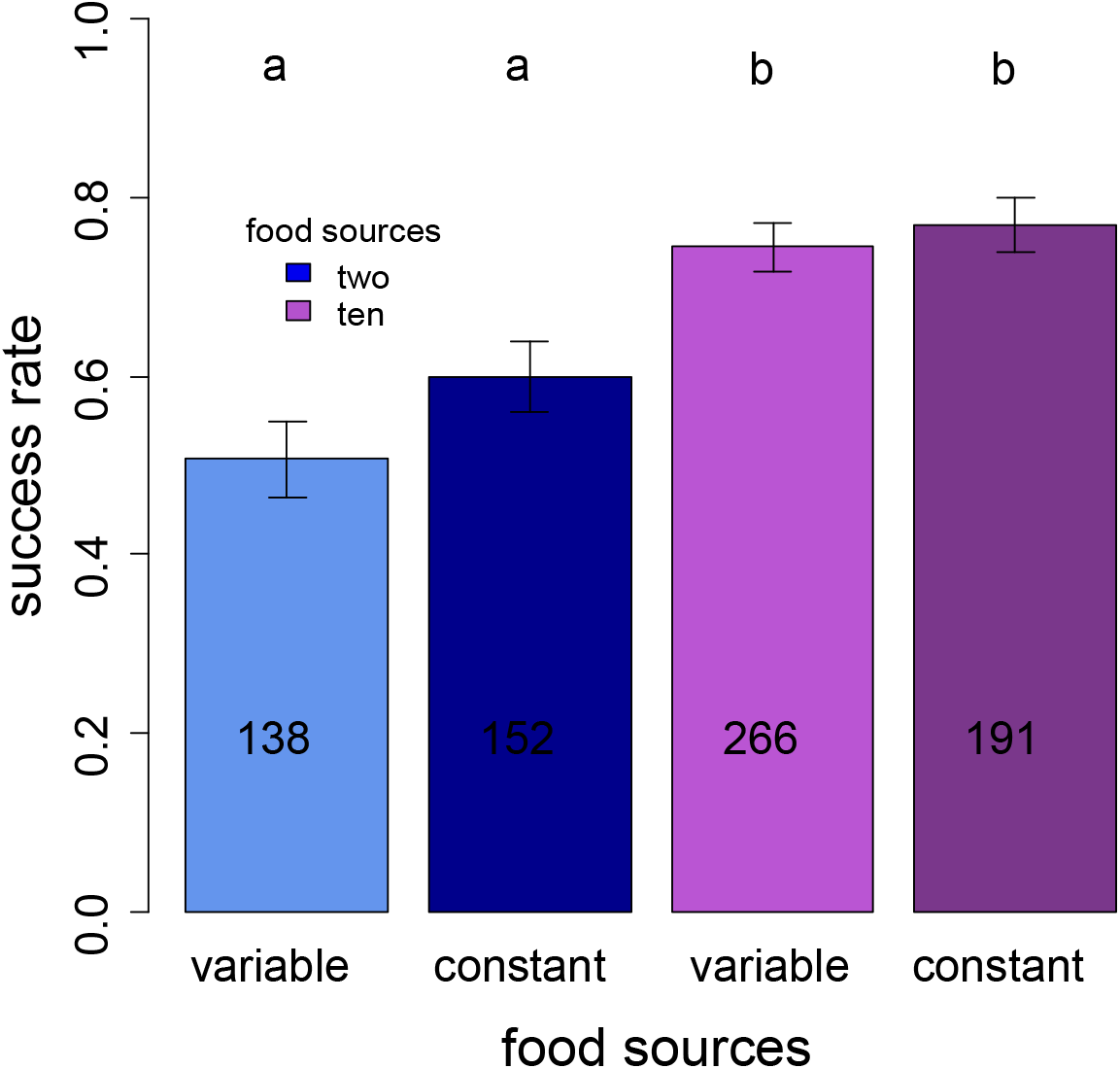
Success rate depending on the food quantity, for two and ten food sources. Numbers in columns represent tandem runs to a food source. Bars show mean ± standard error. Different letters indicate significant differences.

**Fig. 2.**
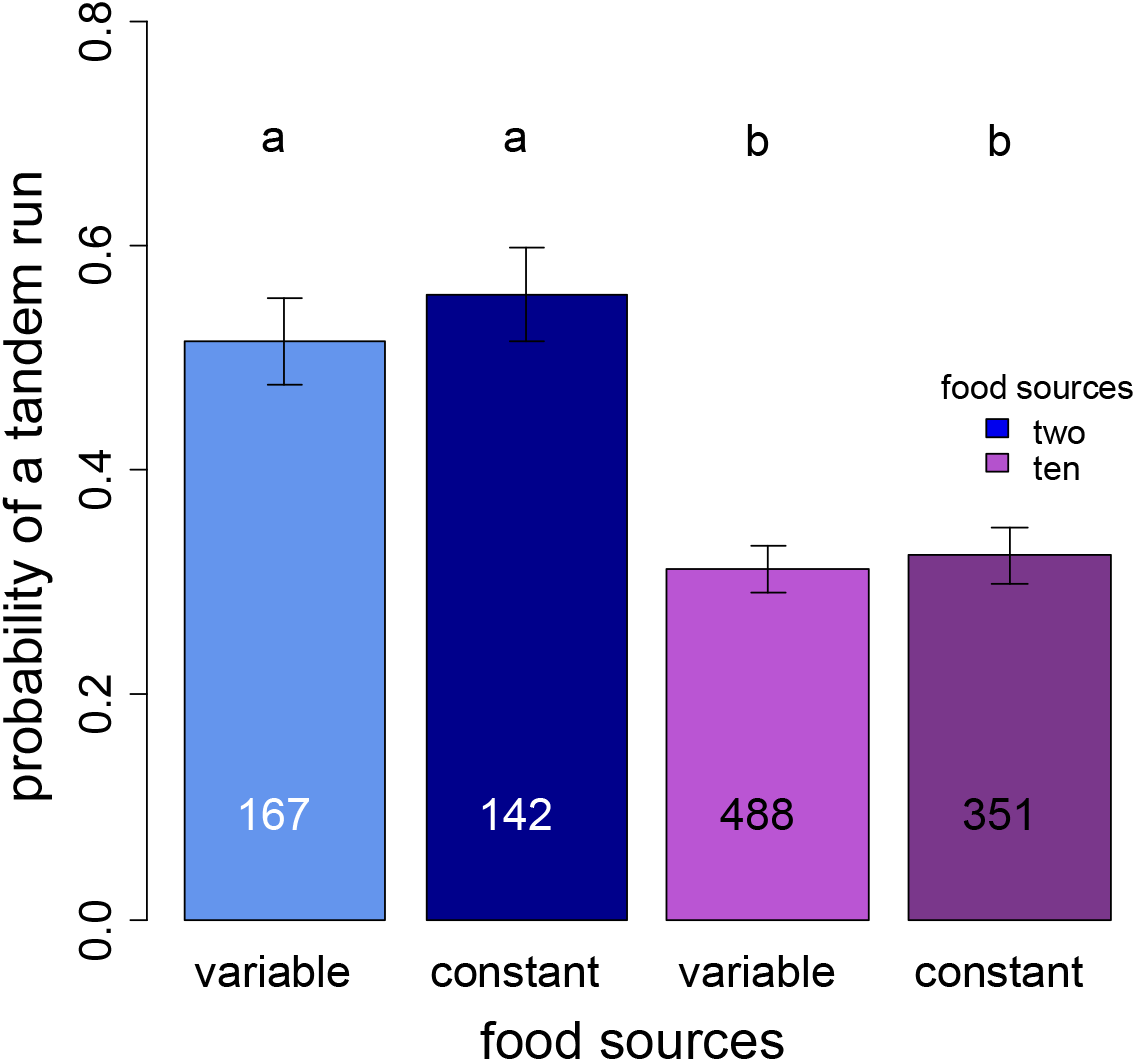
Probability of a tandem run depending on the different setups. Numbers in columns represent the number of individual visits (starting with visit 2).

**Fig. 3.**
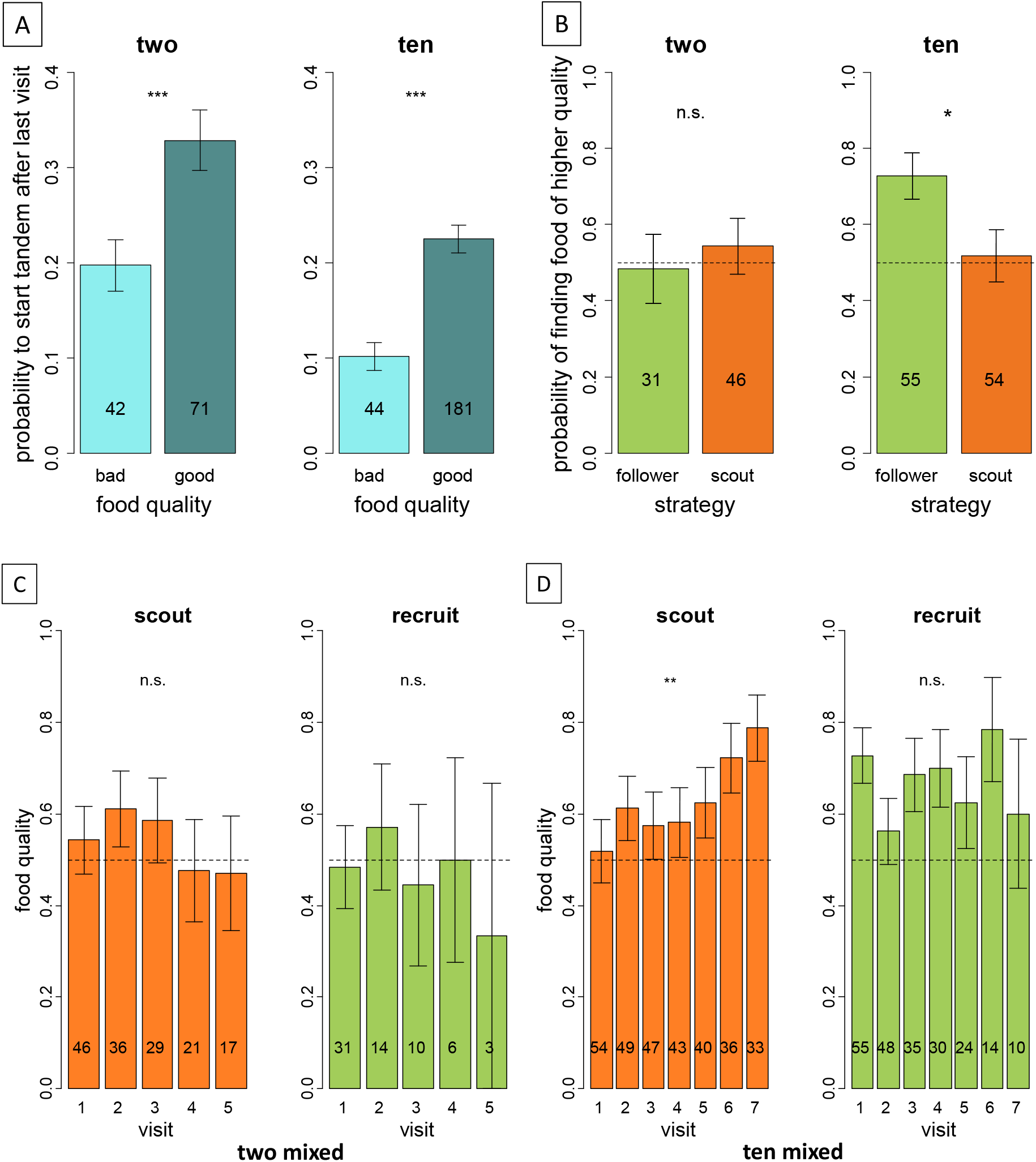
(**A**) Probability to start a tandem run (marked ants) depending on the food quality of the last visit. (**B**) Probability of finding food of a better quality on the first food visit depending on the information-use strategy. Mean food quality depending on the food visits of scouts and recruits for **(C)** few mixed and **(D)** many mixed food sources. Bars show mean ± standard error. **p<0.01, ns: not significant Bars show mean ± standard error. *p<0.05, **p<0.01, ***p<0.001, ns: not significant

### (ii) Quality of food sources discovered by individual and social learners

In contrast to our prediction, scouts and followers found food sources of similar quality when there were only two food sources that differed in quality: 54.4% (25 of 46) of scouts and 48.4% (15 of 31) of followers first found a high-quality food source (Fig. 3B). However, when there were ten food sources, followers were significantly more likely to discover a better food source than scouts (Fig. 3B) (scouts: 51.9% vs. followers: 72.7%) (GLMM: two: z = 0.039, p = 0.969; ten: z = -2.221, p = 0.026). As a result, we found that followers were led to above-average food sources when there were ten food sources (GLMM: z = 2.36, p = 0.018). When there were only two food sources, scouts and original recruits continued to visit food sources of the same, average quality during a trial (Fig. 3C) (GLMM: visits: scouts: z = 0.837, p = 0.403; recruits: z = 1.343, p = 0.179). Strikingly, when there were ten food sources, scouts switched to better food sources over time (Fig. 3D). Recruits continued to visit above average food sources, but without further change in the reward quality over time (GLMM: visits: scouts: z = 2.663, p = 0.008; recruits: z = 0.418, p = 0.676), as they mostly return to the same food source (see below).

### (iii) Probability to switch

When there were two food sources, both scouts and followers visited only 1-1.2 food sources during the 90 minutes, meaning that they largely continued to visit the food source they discovered first. However, switching was more common after visiting a low-quality food source (Fig. 4) (GLMM: food quality: z = 2.258, p = 0.024; strategy: z = 1.082, p = 0.279). With ten food sources, scouts visited on average about 3 different food sources, which was ∼50% more than followers did (Fig. 4A) (GLMM: ten: variable: z = 2.824, p = 0.005; constant: z = 2.835, p = 0.005). Scouts generally switched more often than former followers (Fig. 4B) (GLMM: ten -variable: strategy: z = 2.333, p = 0.020; food quality: z = 0.71, p = 0.478).

**Fig. 4.**
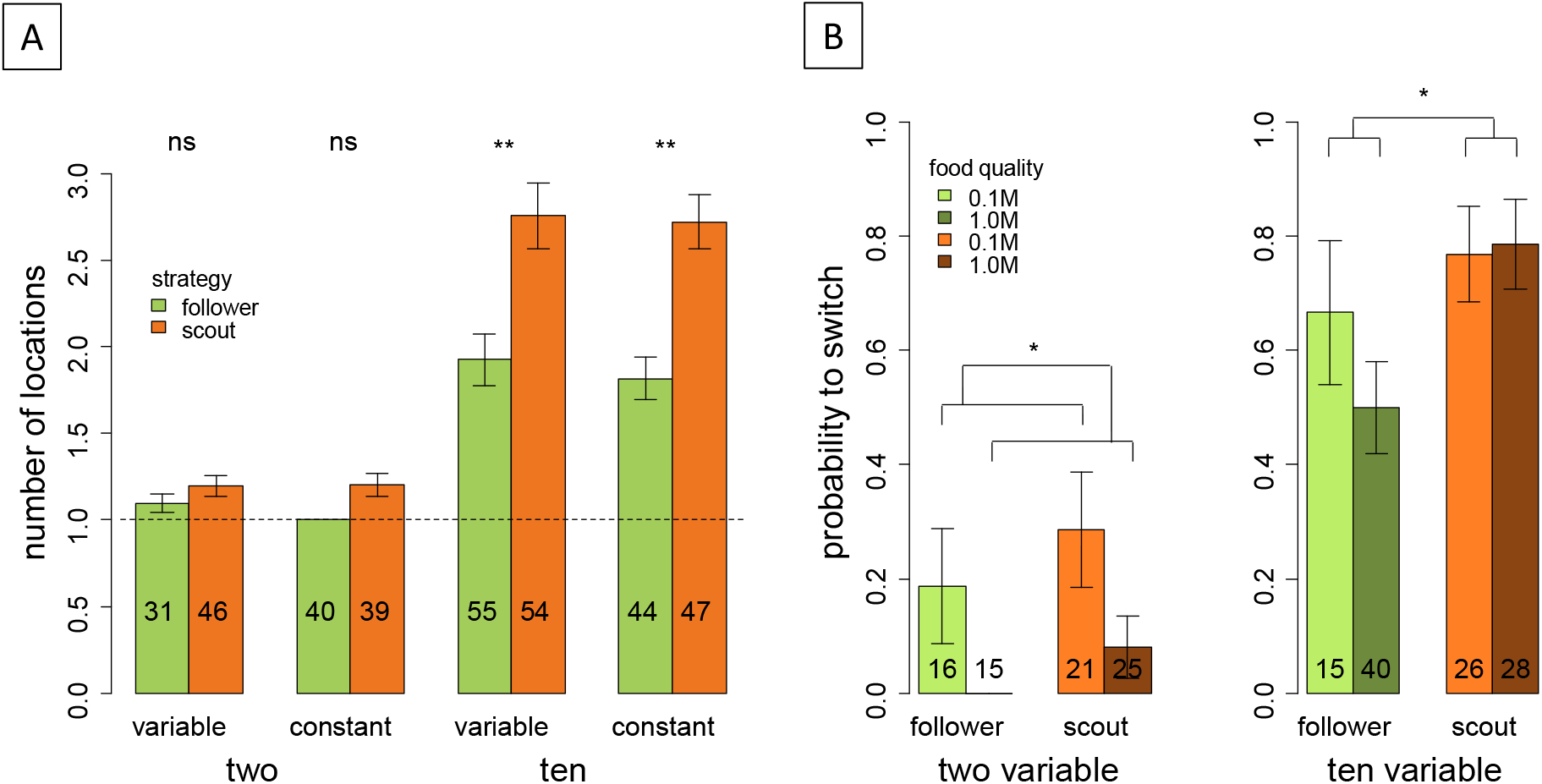
(**A**) The number of food locations scouts and tandem followers visited. (**B**) Probability to switch food source depending on the information-use strategy and the food quality. Number in column match the number of tested ants. Bars show mean ± standard error. *p<0.05, **p<0.01, ***p<0.001, ns: not significant

### (iv) Total foraging activity

We found that scouts performed significantly more trips to food sources than ants that used social learning to discover their first food source, irrespective of the number of food sources that were available (Fig. 5A) (GLMM: follower vs. scout: two: z = 5.405, p < 0.001, ten: z = 8.39, p < 0.001).

**Fig. 5.**
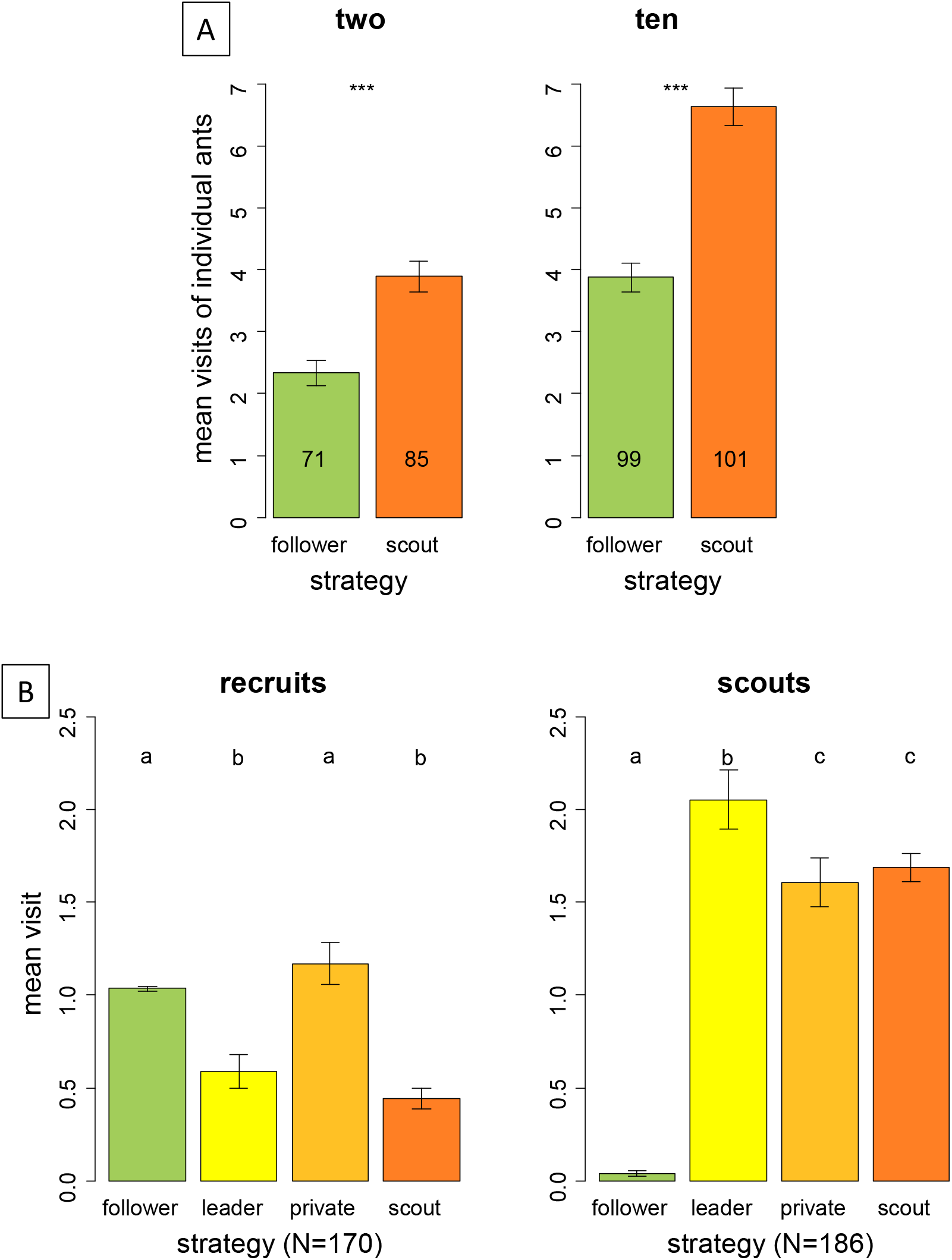
Mean visits of individual ants, depending on the information-use strategy they used to reach a food source. (**A**) Mean visits depending on two and ten food sources. (**B**) Mean visits for individual ants using different information-use strategies for all visits including discovering a food source either as a scout or a recruit. Number in column match the number of ants tested. Bars show mean ± standard error. Different letters indicate significant differences.

When we explored how often scouts and followers followed different strategies after the initial food source discovery (Fig. 5B), we found that former followers usually used private information to return to the same food source (including visits as tandem leaders). They performed fewer individual exploration trips and led fewer tandem runs than former scouts (Table 1). On the other hand, ants that discovered a food source *via* individual learning (scouting) almost never followed tandem runs during the trials (3.8% of 186 cases), but they frequently led them and, therefore, were important providers of social information. They continued to scout and discover new food sources (Fig. 5B, Table 1).

**Table 1:** Information-use strategies of social learners (recruits) and individual learners (scouts). Generalized mixed-effects models (GLMMs) were used to calculate z- and p-values.

| Information-use strategy |  | follower | leader | private | scout |
| --- | --- | --- | --- | --- | --- |
| **Social learner/**<br>**recruit** | follower | - | $z = -4.573$<br>$p < 0.001$ | $z = 1.202$<br>$p = 0.229$ | $z = -6.266$<br>$p < 0.001$ |
| | leader | - | - | $z = 5.687$<br>$p < 0.001$ | $z = -1.908$<br>$p = 0.0564$ |
| | private | - | - | - | $z = -7.295$<br>$p < 0.001$ |
|  | scout | - | - | - | - |
| **Individual learner/**<br>**scout** | follower | - | $z = 10.595$<br>$p < 0.001$ | $z = 9.922$<br>$p < 0.001$ | $z = 10.056$<br>$p < 0.001$ |
| | leader | - | - | $z = -3.206$<br>$p = 0.001$ | $z = -2.600$<br>$p = 0.009$ |
| | private | - | - | - | $z = 0.612$<br>$p = 0.540$ |
|  | scout | - | - | - | - |

### Time until food discovery using social or individual learning

We measured the time tandem followers and scouts needed from leaving the nest until locating their first food source. Overall, scouts needed nearly twice as long as tandem followers to reach the food source (Fig. 6) (LME: scout vs. tandem: two: t = 3.967, p < 0.001; ten: t = 4.737, p < 0.001). Having more food sources available reduced the time that scouts and followers needed to locate a food source (LME: two vs. ten: scout: t = 2.880, p = 0.007; tandem follower: t = 3.848, p < 0.001).

**Fig. 6.**
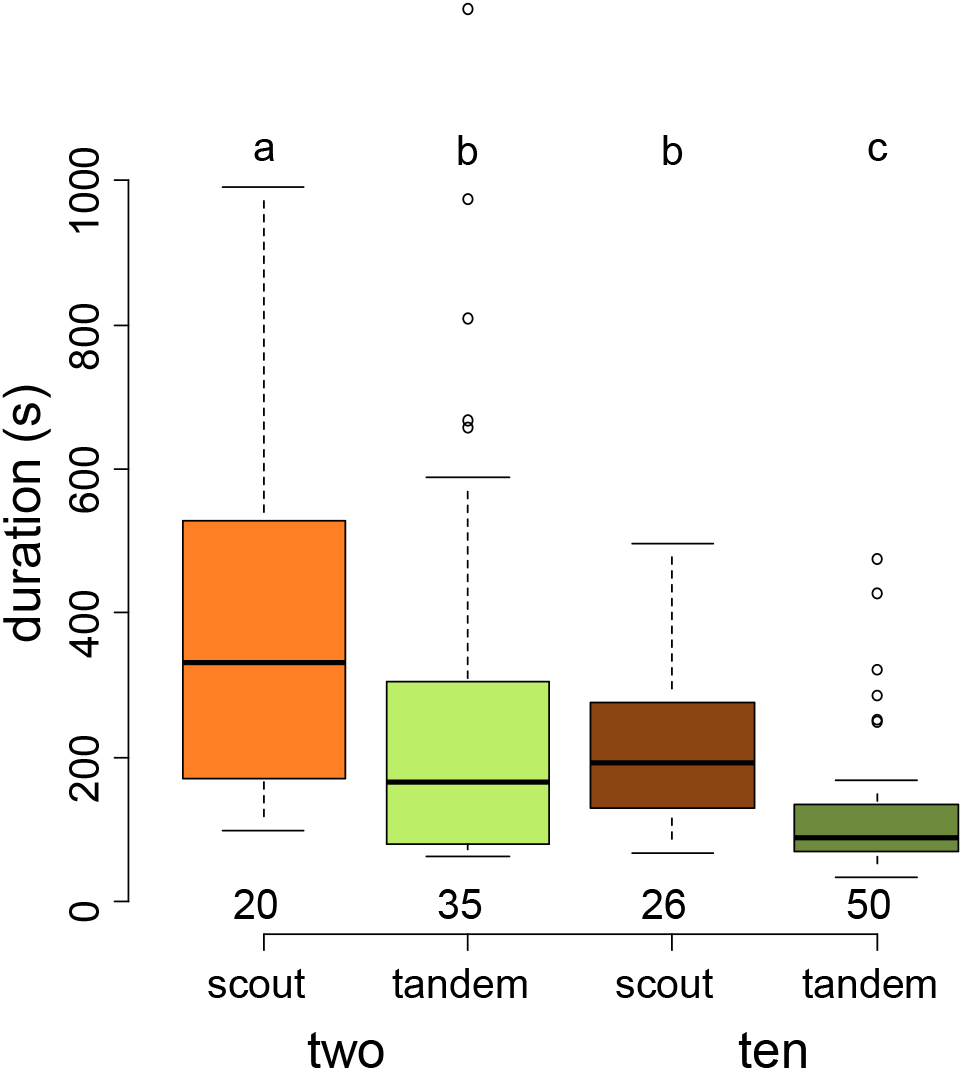
Mean time foragers need to find the first food source. **(A)** Duration of tandem runs and scouts from the nest entrance to a food source. Different letters indicate significant differences.

## Discussion

Simulations predict that social learning leads to better rewards than individual learning because demonstrators filter information for observers, allowing the latter to acquire more successful behavioural strategies (Rendell et al. 2010). Our experimental results show that social learners in *Temnothorax nylanderi* discovered better food sources than individual learners only in some conditions. Only in a rich environment (ten food sources) did social learners locate a better food source than individual learners on their first trip. In such environments, following a tandem run increased the chance of finding a high-quality food source as ants performed more tandem runs to higher quality food sources (Fig. 3). Scouts, on the other hand, initially found food sources of average quality but, they managed to discover food sources of better quality during a trial by following a strategy of food source switching (Fig. 4). In environments where food sources were scarce (two food sources), scouts and recruits discovered food sources of intermediate quality and overall food quality did not change over several visits. The opportunities to find a better food source are much reduced in a poor environment because there are fewer alternatives and food sources are further apart. It is surprising, however, that recruits did not discover better food sources given that more tandem runs were initiated after foragers visited a high-quality food source (Fig. 3) (see also Shaffer et al. 2013). However, the effect of food source quality on tandem runs was less pronounced in a poor environment (Fig. 3A) and tandem runs were more frequently unsuccessful (Fig. 1), which can explain the more similar outcomes for social and individual learners. Our finding that ants were overall more likely to perform tandem runs in a poor foraging environment (Fig. 2) was unexpected, but could be an adaptive response since social information is likely to be more useful to nestmates under these circumstances (see also Dornhaus et al. 2006; Beekman and Lew 2007; Schürch and Grüter 2014; Goy et al. 2021). Similarly, honey bee foragers adjust their dance threshold according to the quality of the foraging environment (von Frisch 1967). Scouts could assess their environment based on the time they needed to locate a food source. More generally, these results suggest that information filtering by demonstrators (Rendell et al. 2010) might not always occur or be effective, e.g. because acquiring information about the payoffs of different behavioural choices is too costly or alternative choices are not available.

One way to improve rewards over time is to abandon poor food sources and switch to better ones. As mentioned, switching by either forager type was rare in our experiments when there were only two food sources. However, switching was more frequent when there were more options, particularly after foragers fed on food of low quality (Fig. 4b). Scouts visited more food sources than tandem followers, suggesting a strategy of actively searching for alternatives, whereas tandem followers mostly used private information to return to the location they were guided to (Fig. 5B). Frequent food source switching not only allows scouts to improve the quality of exploited food sources over time, but also highlights their role as “innovators” that discover new food sources for the colony (von Frisch 1967; Seeley 1983; Liang et al. 2012). Individual learning could be particularly important in an unpredictable and changing environment (Galef and Laland 2005; Kendal et al. 2005; Rendell et al. 2010; Goy et al. 2021).

Our experimental results indicate that there are two types of foragers in *T. nylanderi* that differ in whether they use social or individual learning to discover a food source, similar to the situation in honeybees (Seeley 1983; Liang et al. 2012). Accordingly, Alleman et al. (2019) found that scouts and tandem followers differed in their patterns of brain gene expression during nest migrations (*Temnothorax longispinosus*) and host raids (*T. americanus*). In our trials, scouts were the main providers of social information as they led most of the tandem runs. It is also noteworthy that after the first food source was discovered, the most frequently followed strategy by both scouts and recruits was the use of private information. This is a widespread foraging strategy in social insects (e.g. reviewed in Grüter and Leadbeater 2014; Grüter and Czaczkes 2019). Future research could explore if social and individual learners differ in other aspects, such as their morphology, physiology or age.

Tandem followers needed less time from the nest to the food source than scouts (Fig. 6) (see also Franks and Richardson 2006). On the other hand, we found that social learners performed 60% fewer foraging trips during our experimental trials. This highlights that there are hidden time and opportunity costs to social learning as social learners need to wait for information inside the nest (Dechaume-Moncharmont et al. 2005; Schürch and Grüter 2014). Agent-based simulations that included these waiting times showed that scouts often needed more time than recruits to discover their first food source in environments with two food sources, but they were faster at discovering a food source when there were more and variable food sources (Goy et al. 2021). In the latter situation, it is relatively easy for scouts to find a food source without the help of a nestmate. These findings are in line with empirical observations suggesting that honeybee recruits experience greater time costs than scouts under some conditions (Seeley 1983; Seeley and Visscher 1988).

Communication is often considered to be beneficial, particularly in social insect colonies (but see e.g. Dechaume-Moncharmont et al. 2005; I’Anson Price et al. 2019). However, simulations highlight that there is often a narrow parameter space that favours colonies to use recruitment communication, especially when species have small colony sizes (Goy et al. 2021). This is consistent with the finding that ants and stingless bees with small colony sizes often forage solitarily (Maschwitz et al. 1974; Jessen and Maschwitz 1986; Beckers et al. 1989; Grüter 2020) and could explain why ants like *Diacamma, Neoponera or Paltothyreus*, bumblebees and many stingless bees do not share information about food source locations with nestmates, even though some of these species use communication during colony emigrations where all colony members need to relocate to the same location. One potentially important factor that was not explored in our study is the role of foraging competition. In highly competitive environments, recruitment communication allows colonies to build up a critical mass of foragers to defend and exploit a food source that would otherwise be lost to competitors (Hrncir & Maia-Silva 2013; Glaser et al. 2021). Detailed natural history information about the foraging ecology of a species is often necessary to understand why foragers do or do not communicate about food source locations.

## Supporting information

Raw data

## Acknowledgement

We thank Julia B. Saltz, Erol Akcay and the members of the Foitzik lab for feedback on the study. S.M.G. and C.G. were funded by the Deutsche Forschungsgemeinschaft (DFG) (GR 4986/1-1).

## Data availability statement

Raw data are available as an online supplement.

## Tables

**Fig. S1.**
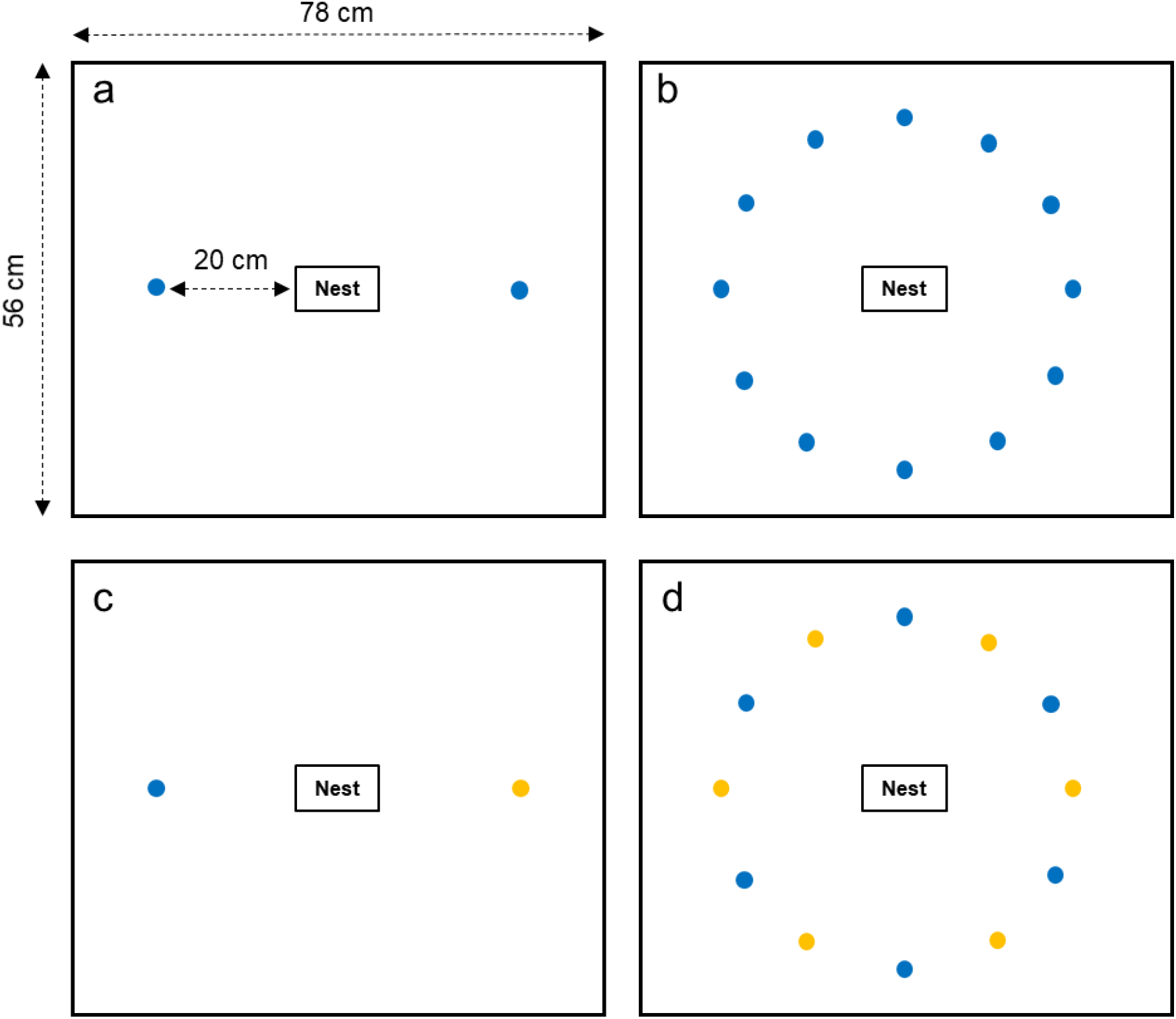
Schematic illustration of the four setups (a-d) and the dimensions of the foraging box and food source distances. Differently colored dots represent high- and low-quality sucrose solution sources (blue and orange, respectively).

